# *Trichinella spiralis* secretes abundant unencapsulated small RNAs with potential effects on host gene expression

**DOI:** 10.1101/2020.02.09.940601

**Authors:** Peter J. Taylor, Jana Hagen, Farid N. Faruqu, Khuloud T. Al-Jamal, Bonnie Quigley, Morgan Beeby, Murray E. Selkirk, Peter Sarkies

## Abstract

Many organisms, including parasitic nematodes, secrete small RNAs into the extracellular environment largely encapsulated within small vesicles. Parasite secreted material often contains microRNAs (miRNAs), raising the possibility that they might contribute to pathology by regulating host genes in target cells. Here we characterise material from the parasitic nematode *Trichinella spiralis* at two different life stages. We show that adult *T. spiralis*, which inhabit intestinal mucosa, secrete miRNAs within vesicles. Unexpectedly however, *T. spiralis* muscle stage larvae (MSL), which live intracellularly within skeletal muscle cells, secrete miRNAs that appear not to be encapsulated. Notably, secreted miRNAs include a homologue of mammalian miRNA-31, which has an important role in muscle development. Our work therefore suggests a new potential mechanism of RNA secretion with implications for the pathology of *T. spiralis* infection.

## Introduction

It is becoming increasingly clear that a variety of RNA molecules have a life outside the cell (Sarkies and Miska, 2014). RNA has been found in many different extracellular fluids (Liu et al., 2019). Often, RNA molecules are encapsulated in membrane bound extracellular vesicles (EVs) such as exosomes and microvesicles, which are presumed to be required to protect them from nuclease digestion. Of the categories of RNA observed, one of the most abundant appears to be small (18-30 nt) RNAs, including microRNAs (miRNAs) which, within cells, are critical regulators of gene expression (Bartel, 2018). Extracellular miRNAs are often represented in different proportions to miRNAs within the cells from which they originate, indicating that specific selection of certain miRNAs for export may occur (Turchinovich et al., 2016).

Whilst the existence of extracellular miRNAs is clear, most aspects of their life cycle are still poorly understood. It is not clear how specific miRNAs are targeted for export, which cytoplasmic proteins are responsible for sorting miRNAs into vesicles, and the protein factors that bind miRNAs outside cells (Sarkies and Miska, 2014). There is also little understanding of the function of extracellular miRNAs. In principle, miRNAs might be taken up by target cells and act to regulate gene expression, thus contributing to cell-to-cell communication (Turchinovich et al., 2016). However, significant challenges remain for this model. The density of miRNAs within EVs is somewhat low, meaning that high numbers might have to be delivered to target cells. Moreover, there is limited evidence that extracellular miRNAs are bound to the Argonaute proteins that are required for their operation (Gibbings et al., 2009), and it is thus unclear how they would be able to function in host gene expression pathways.

Recently, parasitic nematodes have emerged as an attractive system in which to study the potential roles of extracellular miRNAs (Buck et al., 2014; Coakley et al., 2015). Parasitic nematodes secrete abundant material in order to develop and complete their life cycle within their hosts, often involving manipulation of host immunity or physiology (Coakley et al., 2016). The secreted material from many nematode species is enriched for small RNAs including miRNAs (Buck et al., 2014). Communication with the host via secreted miRNAs might be a key mechanism in parasitic infection, as the high conservation of miRNA sequences and mechanism across animals (Bartel, 2018) would facilitate their incorporation into host gene expression pathways. Additionally, the crucial importance of many miRNA-target interactions in development and cellular homeostasis (Gebert and MacRae, 2019) would make it difficult for host cells to evolve resistance mechanisms. Many examples have been cited whereby exposure to parasite-derived EVs led to modified immune responses in mammalian cells (Coakley et al., 2015). Exosomes secreted by *Heligmosomoides polygyrus* containing miRNAs and other small RNAs were taken up by murine epithelial cells in vitro resulting in altered gene expression, and intranasal delivery of parasite exosomes modified the pulmonary immune response to a co-administered fungal extract (Buck et al., 2014). Nevertheless, there is still little concrete evidence that these pathways occur *in vivo*, and particularly challenging is the question of whether secreted vesicles can be taken up by target cells at a high enough rate to ensure delivery of miRNAs at robust levels, although parasite-derived miRNAs were recently identified in macrophages from the pleural/peritoneal cavity of Mongolian jirds infected with the filarial nematode *Litomosoides sigmodontis* (Quintana et al., 2019).

Here we describe the nematode *Trichinella spiralis* as a new model to study extracellular miRNAs. *T. spiralis* is a parasite that infects many mammalian species including humans. Its life cycle is unusual amongst parasitic nematodes because it has both intracellular and extracellular stages. In the intestinal phase, infective larvae penetrate and migrate through epithelial cell sheets (ManWarren et al., 1997; Wright, 1979), and following development to adult worms, release first stage larvae which migrate via the lymphatics and vascular system to skeletal muscle, where they invade myofibres and develop to form a specialised intracellular niche known as the nurse cell. Interestingly, nurse cell formation involves perturbation of the normal gene expression programme of muscle cells, which results in cell cycle re-entry, arrest at apparent G2/M phase and downregulation of several key markers of differentiated muscle (Jasmer, 1993). How the parasite influences gene expression and subsequent alterations in skeletal muscle phenotype is poorly understood.

We reasoned that the existence of *Trichinella* in such radically different environments provided an interesting opportunity to study the potential role of RNA in extracellular communication. In particular, release of RNA from intracellular parasites might be expected to facilitate a high local concentration of parasite miRNAs which otherwise would be difficult to achieve through uptake of vesicles by target cells. We therefore isolated secreted material from adult and larval *T. spiralis* and investigated the RNA content. Whilst adult *T. spiralis* releases miRNAs protected from RNase digestion, larvae isolated from muscle cells secrete unprotected small RNAs, suggesting they might be released directly into target cells. Amongst the miRNAs secreted by muscle stage larvae, we identified a homologue of the mammalian miR-31, which plays a key role in development and regeneration of skeletal muscle. Our results highlight a new paradigm for secreted miRNAs without vesicular protection, and suggest a novel mechanism for how intracellular *T. spiralis* may affect mammalian muscle gene expression.

## Materials and Methods

### Parasite isolation and culture

This study was licensed by and performed under the UK Home Office Animals (Scientific Procedures) Act Personal Project Licence number 70/8193: ‘Immunomodulation by helminth parasites’. Adult parasites were recovered from Sprague-Dawley rats 6-7 days post-infection by sedimentation in a Baermann apparatus containing segments of small intestine, and muscle stage (infective) larvae were recovered from digested muscle 2 months post-infection as previously described (Arden et al., 1997). Parasites were cultured in serum-free medium for up to 72 hrs as described (Arden et al., 1997), secreted products collected daily, centrifuged at 2,000 x g for 10 min, supernatants cleared through 0.2 µm filters and pooled.

### Isolation and analysis of extracellular vesicles

Culture medium was first centrifuged at 10,000 x g for 30 min to clear cell debris, apoptotic bodies and large vesicles. The supernatant was then centrifuged at 100,000 x g for 90 min in polyallomer tubes at 4°C in a SW40 rotor, the pellet washed in PBS and recentrifuged under the same conditions. For Nanoparticle Tracking Analysis (NTA), the pellet was resuspended in 400 µl PBS and analysed as previously described (Faruqu et al., 2018). For electron microscopy, the pellet was fixed in 2% paraformaldehyde, adsorbed onto copper EM grids, washed with PBS, stained with uranyl acetate and viewed in a Tecnai T12 Spirit Electron Microscope with images captured on a TVIPS TemCam-F216 CCD camera.

### RNase protection assay

Parasite secreted products were passed through a 0.2 µm filter and concentrated in 3 kDa molecular weight cutoff vivaspin columns, washed once in PBS, then 3x in RNase buffer (PBS, 5 mM EDTA, 300 mM NaCl, 10 mM Tris–Cl pH 7.5). Aliquots (150 µl) of concentrated secreted products were digested with different concentrations of RNase A/RNase T1 cocktail (2 µg RNase A/5 U RNase T1, 40 ng RNase A/0.1 U RNase T1, no enzyme control) for 1 hr at 37°C, and digestion terminated by addition of 700 µl Trizol. RNA was isolated from each sample and resolved on a 2100 Bioanalyser using the Agilent small RNA kit in accordance with instructions.

### Small RNA sequencing

RNA was isolated from parasite secreted products using TRIzol, and libraries prepared using the Illumina TruSeq small RNA preparation kit. Sensitivity to RNase was again assessed by digesting some RNA samples with 4 µg ml^-1^ RNase A/10 U ml^-1^ RNase T1 at 37°C for 1 hr prior to library preparation. Sequencing was performed by the MRC London Institute of Medical Sciences High Throughput Sequencing facility using a Hiseq 2000. For data analysis, adapters were removed using fastx-trimmer, and files collapsed using fastx-collapser. First nucleotide and length for each sequence were extracted using a custom Perl script. To identify miRNAs, miRDeep2 annotation of *T. spiralis* performed previously (Sarkies et al., 2015) was used, and mature miRNA sequences were identified in each library by sequence matching using a custom Perl script. Blast was used to examine sequence files for any matches to Y-RNA. Potential targets in mouse 3’UTRs were assessed by searching for complementary sequences to the seed sequence (nucleotides 2-8) within the 3’UTRs of mouse RNAs, downloaded from Ensembl. All figures were prepared in R.

## Results

In order to characterise the RNA content of secreted material from *T. spiralis*, we isolated secreted material from adult worms and muscle stage larvae (MSL). We treated secreted material with different concentrations of exonuclease to test whether the extracellular RNA was encapsulated in EVs or otherwise protected, then extracted RNA and analysed size and quantity using a bioanalyzer. We recovered abundant small RNAs from both adults and larvae. Adult small RNA was largely unaffected by RNase treatment, indicating that it was protected from exonuclease digestion. In contrast, the majority of small RNA from larvae was degraded by the same treatment (Figure 1A-F).

**Fig. 1.**
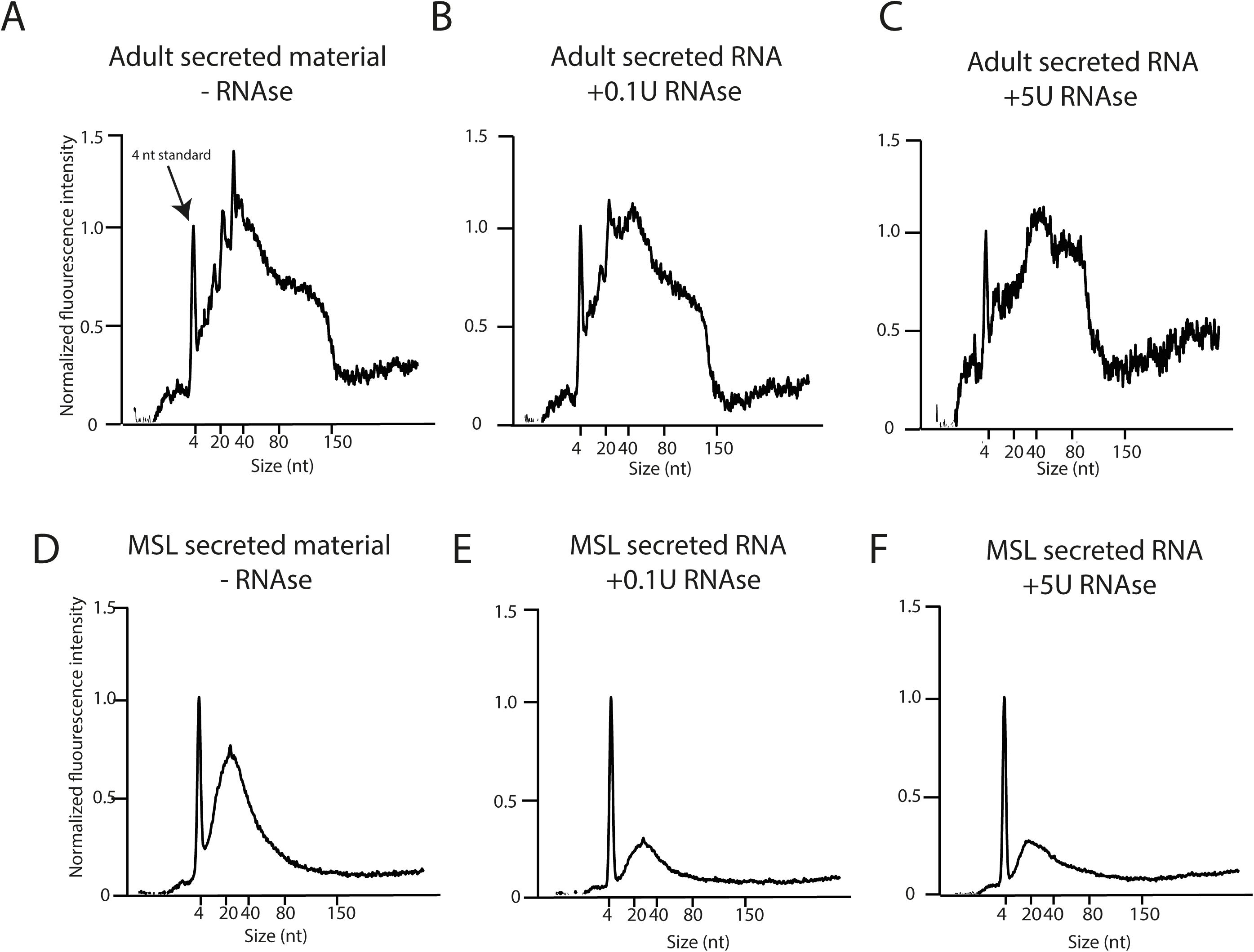
Profile of RNA in secreted material. Bioanalyzer traces showing small RNA profiles from secreted material extracted from Adult (A-C) and Muscle Stage Larvae (MSL; D-F) after exposure to different concentrations of exonuclease.

The lack of protection of small RNAs could reflect absence of vesicles in secreted material. We therefore used a standard protocol to isolate extracellular vesicles from MSL secreted products by ultracentrifugation, and examined the pelleted material by Transmission Electron Microscopy (TEM), which revealed vesicle-like structures up to 150 nm in diameter (Figure 2A). To confirm this, we used Nanoparticle Tracking Analysis (NTA) to profile secreted material from MSL. The results showed a disperse profile of particle size consistent with TEM images, confirming the presence of EVs with many in the size range typical of exosomes (Figure 2B), with a similar profile observed by NTA of secreted material from adult worms (Figure 2C). Taken together with the lack of exonuclease protection, this indicates that whilst MSL do secrete EVs, the majority of secreted small RNAs do not appear to be contained within these structures. Quantitation of the number of vesicles secreted by each life stage indicated that over the 72 hr period of maintenance *in vitro*, adult *T. spiralis* secreted on average 3.15 ×10^5^ vesicles per parasite per 24 hrs, in comparison to 1.26 × 10^5^ vesicles per parasite per 24 hrs for MSL. Given that adult worms are estimated to be between 1.4 to 4 x the length of larvae, this is indicative of a broadly similar rate of vesicle production in terms of body mass.

**Fig. 2.**
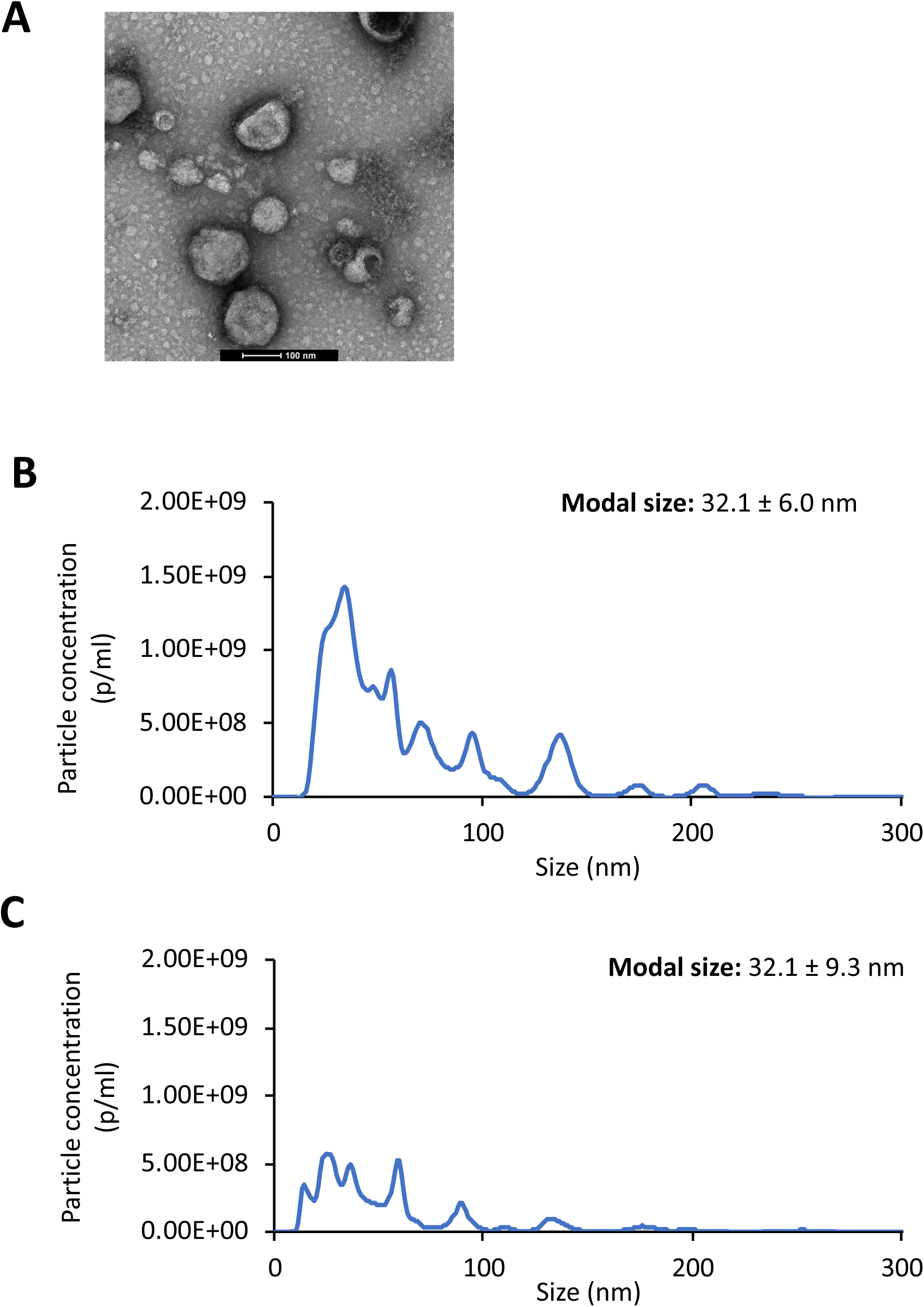
Characterisation of extracellular vesicles (EVs) secreted by parasites at different life stages. A. Transmission electron microscopy (TEM) images of EVs derived from MSL parasites. B, C. Histograms showing size distribution of EVs derived from muscle stage larvae (MSL) and adult parasites, respectively, measured by nanoparticle tracking analysis (NTA). Modal size was considered as the representative size of the vesicle population isolated from conditioned culture medium. Values were expressed as mean ± SD, where n=4 measurements per sample.

We next characterised parasite small RNAs by high-throughput sequencing. Material extracted from both adult and MSL somatic tissue showed a profile peaking at 23 nucleotides in length with a substantial fraction of 24 and 25 nucleotide small RNAs. This is consistent with our previous demonstration that *T. spiralis* produces abundant endo-siRNAs with 23-25 nucleotide length that map to transposable elements (Sarkies et al., 2015). In contrast, secreted RNAs from both larvae and adult worms peaked at 23 nucleotides in length (Figure 3A and D), and thus the small RNAs derived from transposable elements are most likely depleted from secreted material. We then assessed whether small RNAs in secreted material were protected from exonuclease digestion. The profile of secreted small RNAs from adult worms was similar with and without exonuclease treatment, however in larvae the majority of small RNAs were removed by exonuclease treatment under the same conditions, suggesting that most small RNAs in MSL secreted material are unprotected (Figure 3B, C, E and F). Consistent with this observation, small RNAs from purified EVs from MSL showed a flat profile, whilst small RNAs in the supernatant were enriched for small RNAs with a size of ∼22 nucleotides (Figure 3G and H). Of note, in contrast to other studies on parasitic nematode secreted material (Buck et al., 2014) we did not detect Y-RNAs in the secreted material from *T. spiralis* or in the material isolated from whole worms. However, the Y-RNA family has yet to be described in *T. spiralis*, potentially due to the rapid divergence in sequence over evolutionary distance (Boria et al., 2010), and thus we cannot exclude the presence of Y-RNA in secreted material. Again in contrast to earlier studies on parasitic nematodes(Buck et al., 2014; Chow et al., 2019), 22G-RNAs were not found in secreted material, as these evolved in Chromadorea nematodes and are not found in *T. spiralis* (Sarkies et al., 2015) miRNAs are an important component of RNA secreted by parasitic nematodes (Buck et al., 2014; Quintana et al., 2019). We therefore mapped small RNAs from secreted material to our previous annotations of miRNAs in *T. spiralis* (Sarkies et al., 2015). Secreted products contained approximately 15% miRNA in both larvae and adults, around 2-fold depleted relative to the whole worm. This fraction was similar for adult material treated with exonuclease, suggesting that most miRNAs secreted by adult worms are protected from exonuclease activity, in contrast to miRNAs in MSL secreted products (Figure 3I). We found several specific miRNAs at high levels in secreted products. Most miRNAs from MSL were completely removed by exonuclease treatment, whereas miRNAs secreted by adult parasites were protected from digestion. Many miRNAs showed differential abundance in secretions relative to total worm tissue (Figure 4). The majority of miRNAs were depleted in secreted products whilst a small number were enriched (Figure 5A), suggesting some selectivity in which miRNAs are secreted relative to those retained within cells. Notably, the correlation between enrichment in MSL and adult secreted material was weak (Figure 5B), supporting different mechanisms of secretion between the two stages.

**Fig. 3.**
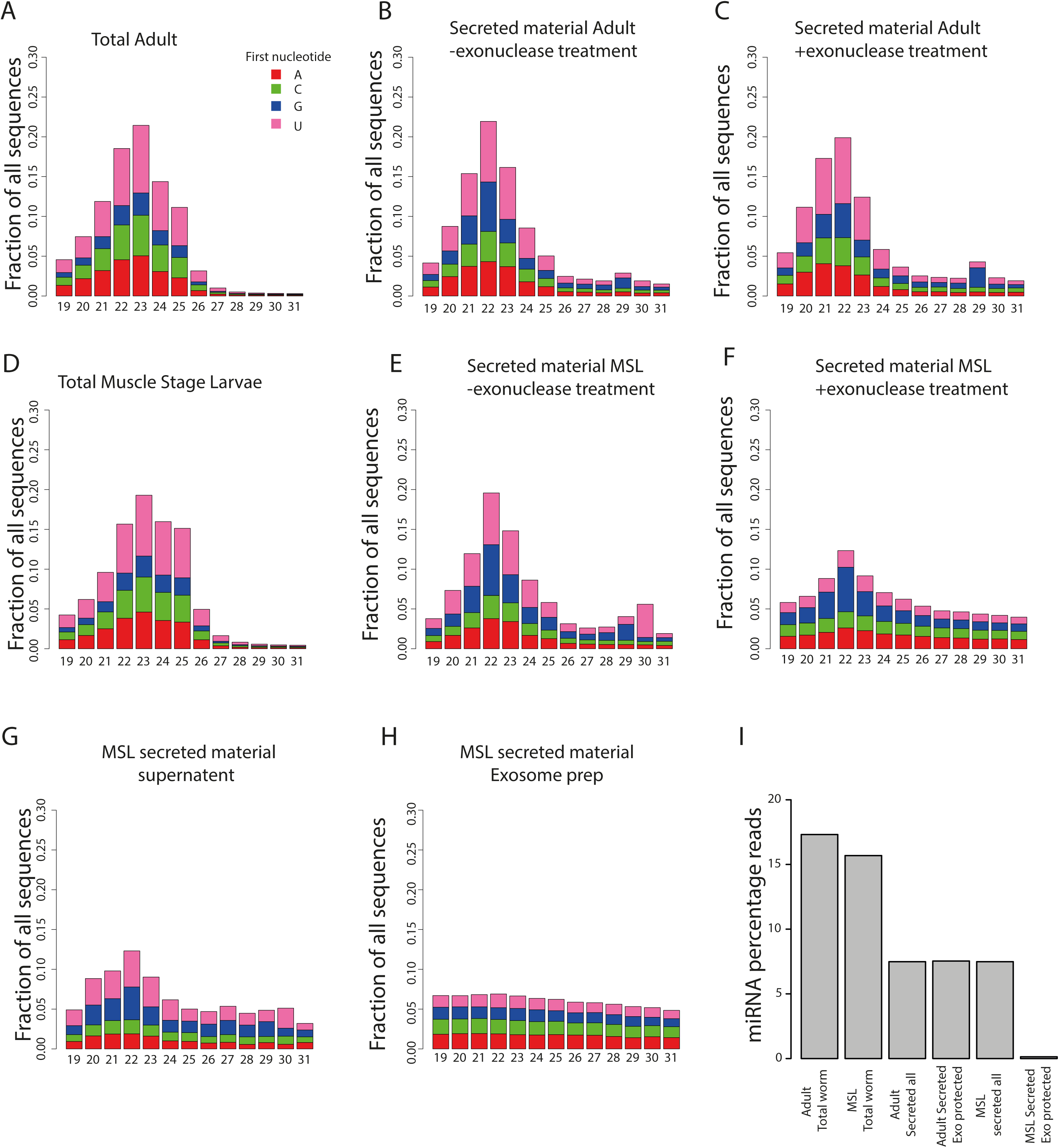
Characterisation of small RNAs secreted by *T. spiralis* by high-throughput sequencing. A-C. The length and first nucleotide of small RNA sequences in adult secreted material either with or without exonuclease treatment compared to total worm material. D-E. The length and first nucleotide of small RNA sequences in MSL secreted material either with or without exonuclease treatment compared to total worm material. G, H. Supernatant (unencapsulated) or pellet (exosomes) after ultracentrifugation of MSL secreted material. I. Fraction of reads mapping to miRNAs in total worm extracts compared to secreted material.

**Fig. 4.**
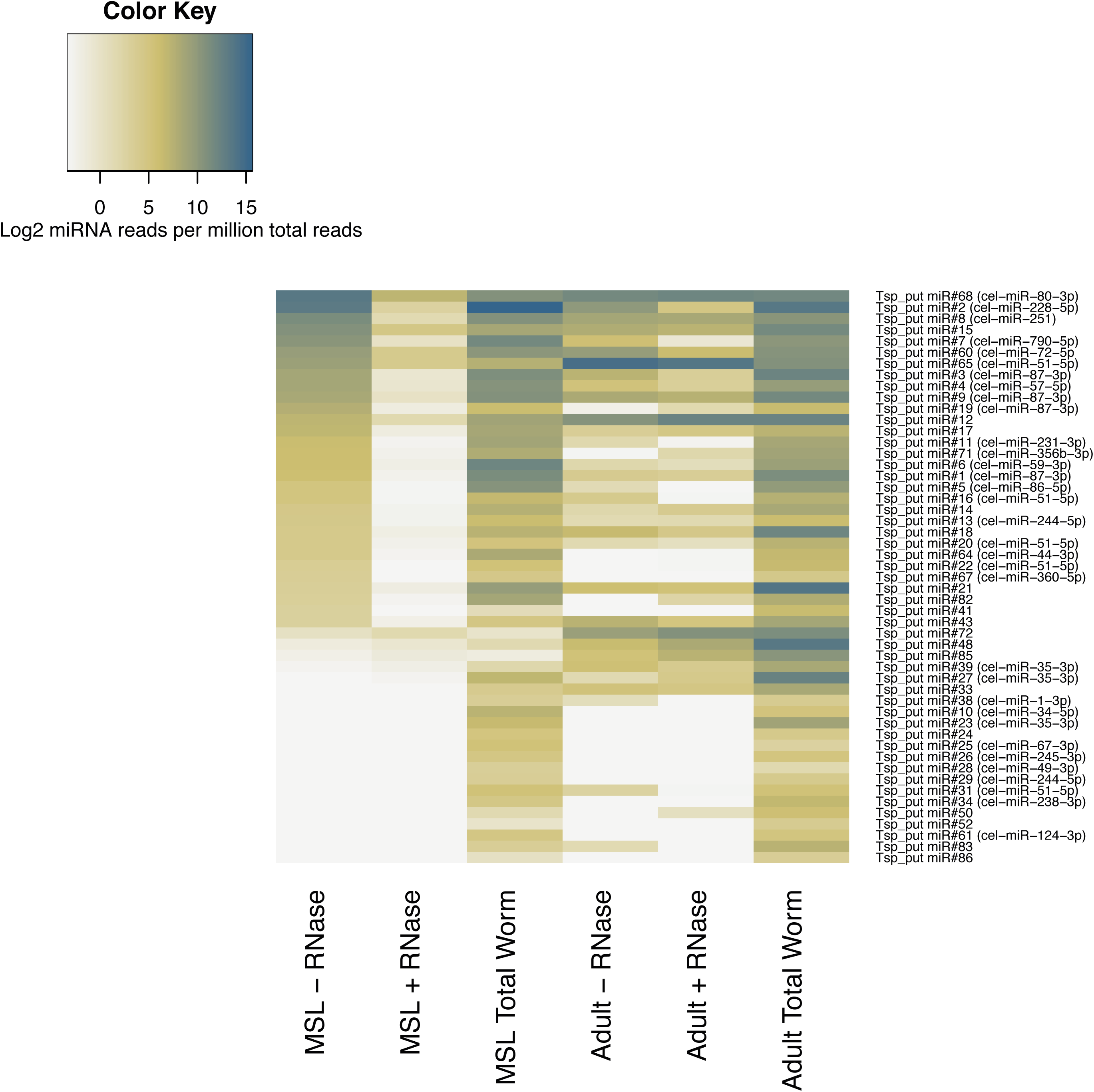
Characterisation of miRNAs in secreted material. A. Heatmap indicating the normalized abundances of miRNAs. miRNAs are sorted in descending order of abundance in MSL secreted material.

**Fig. 5.**
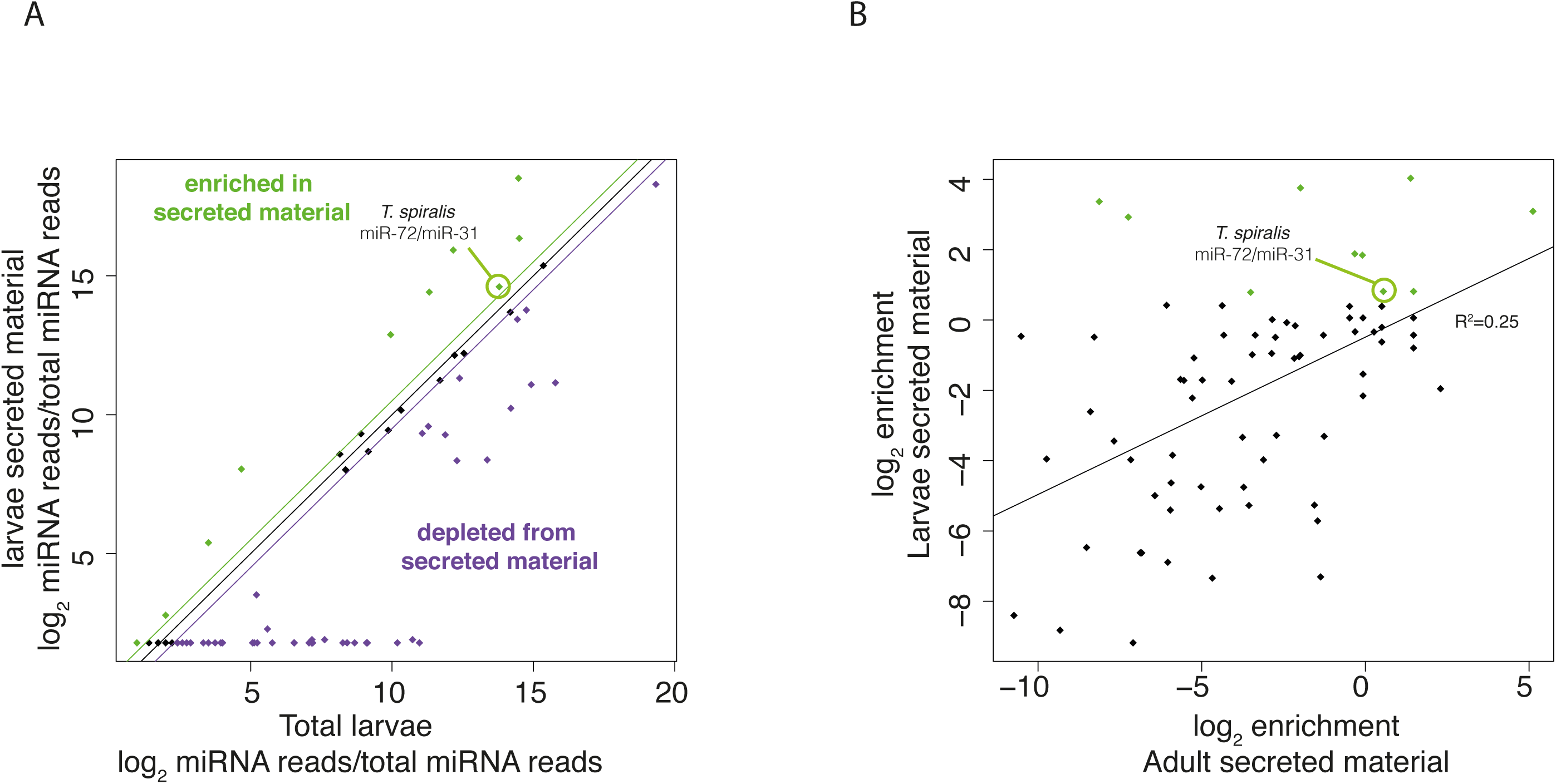
A. Scatter plot indicating the relationship between abundance in total worm and abundance in secreted material. The black line indicates equal abundance, the green line indicates a ∼40% enrichment (log2 0.5) and purple a ∼40% depletion. B. Scatter plot showing correlation between enrichment in adult secreted material and larval secreted material. The line of best fit according to a linear model is shown along with the correlation coefficient.

To investigate the possible consequences of miRNA secretion by *T. spiralis* larvae we identified miRNAs >1.4-fold enriched in the secreted material (Figure 5A; Table 1). Interestingly these included a homologue of mammalian miR-31, previously annotated as a homologue of *C. elegans* miR-72 (Figure 5, Table 1). This was notable because miR-31 has a well-documented role in muscle development by regulating translation of the myogenic determination gene *Myf5*. In particular, miR-31 is transcribed at high levels in quiescent satellite cells, and is sequestered along with *Myf5* RNA in messenger ribonucleoprotein (mRNP) granules. Following activation of satellite cells during development or after injury, mRNP granules are dissociated, miR-31 is reduced, and *Myf5* mRNA is released from repression allowing myogenesis to proceed (Crist et al., 2012). Some enriched miRNAs showed no homology to mammalian miRNAs but nevertheless were predicted to target sites in mammalian 3’UTRs (Table 1) indicating that exposure to these miRNAs may have effects on gene expression in infected cells.

**Table 1.**
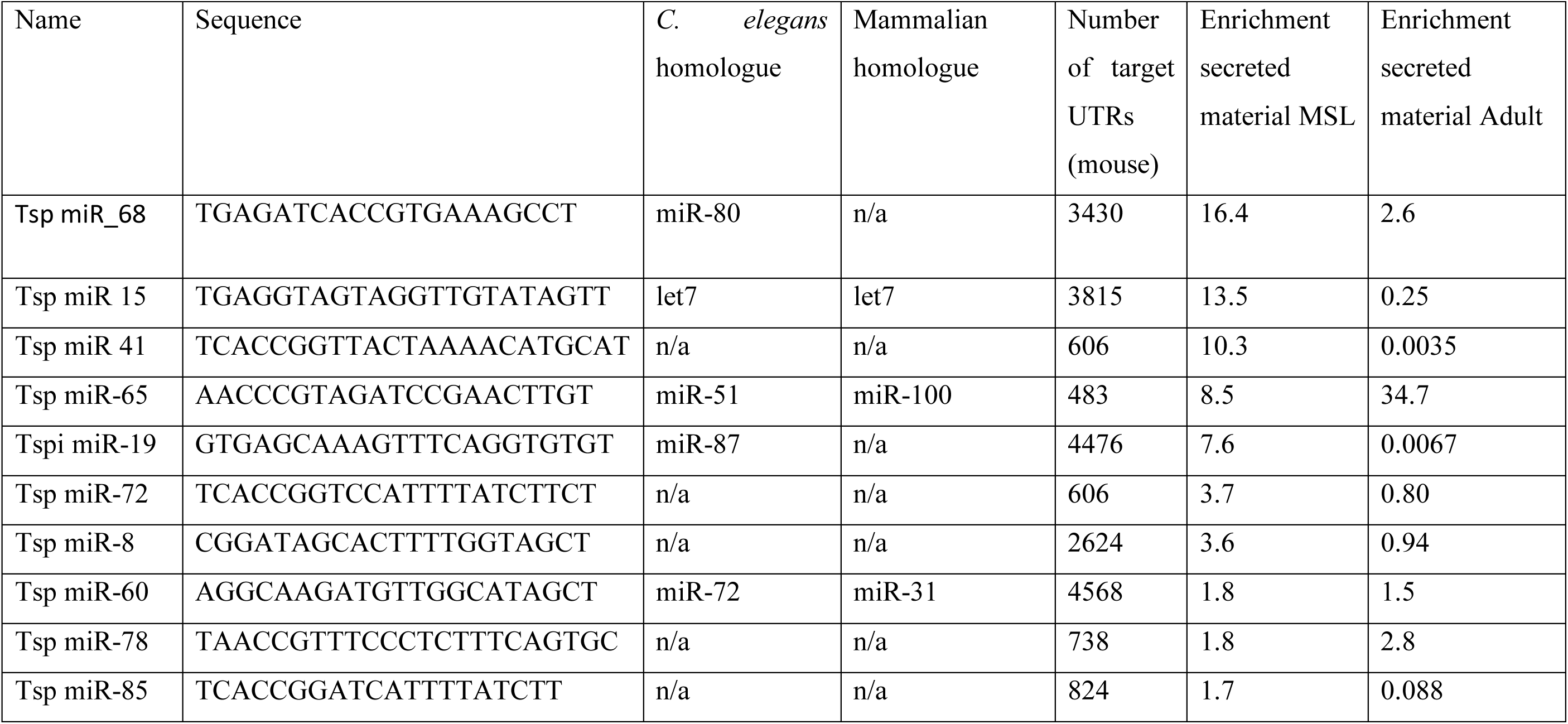
miRNAs found in secreted material. The sequence, homology, and number of potential targets in mammalian 3’UTRs are shown.

## Discussion

Small RNAs encapsulated in EVs are secreted from many cell types, and there is growing evidence that this is a common feature of all the major classes of helminth parasites (Tritten and Geary, 2018). Small RNAs secreted by the parasitic nematodes *Heligmosomoides polygyrus* and *Litomosoides sigmodontis* are resistant to degradation by RNase, but sensitive in the presence of Triton X-100, indicative of encapsulation in membrane-bound vesicles (Buck et al., 2014; Quintana et al., 2019). The number of exosome-sized particles released by *L. sigmodontis* as determined by NTA was very low in comparison to *H. polygyrus*, and sensitivity to RNase was also conferred by exposure to proteinase K, suggesting that extracellular RNAs may also be stabilised via interaction with proteins (Quintana et al., 2019). Analysis of small RNAs secreted by larval *Schistosoma mansoni* identified a comparable abundance of miRNAs and tsRNAs in EV-enriched and EV-depleted fractions, again suggesting extracellular existence outside vesicles and possible stabilisation by proteins (Nowacki et al., 2015).

Our results indicate that adult *T. spiralis* secrete small RNAs that are resistant to exonuclease digestion and most likely encapsulated within vesicles, but that those secreted by MSL appear to be unencapsulated. The authors of the study on *S. mansoni* suggested that small RNAs might be secreted in vesicles which were highly labile, lysing during *in vitro* culture to release their contents (Nowacki et al., 2015). Whilst that is possible, our current data were obtained via culture of two different stages of *T. spiralis* and subsequent processing of material under identical conditions, and intrinsic lability of vesicles from muscle stage larvae alone appears unlikely. In addition, the few miRNAs that are resistant to exonuclease treatment in larval secreted material do not show similar abundance to those that are sensitive (Figure 4, Figure 5B). We suggest that *T. spiralis* larvae secrete small RNAs via an alternative mechanism, potentially involving release directly or accompanied by a chaperone (Figure 6), but this is highly speculative and remains a question for future work. Recent studies have discovered a secreted Argonaute that binds to siRNAs, though not miRNAs, in secreted material from parasitic nematodes within exosomes (Chow et al., 2019). Whether a similar process is implicated in *T. spiralis* secretion of unencapsulated small RNAs is still to be determined. Nevertheless, the observation that not all miRNAs found in secreted material from other helminth parasites appear to be encapsulated within vesicles (Buck et al., 2014; Nowacki et al., 2015) suggests that *T. spiralis* may be an extreme example of a process found in other species..

**Fig. 6.**
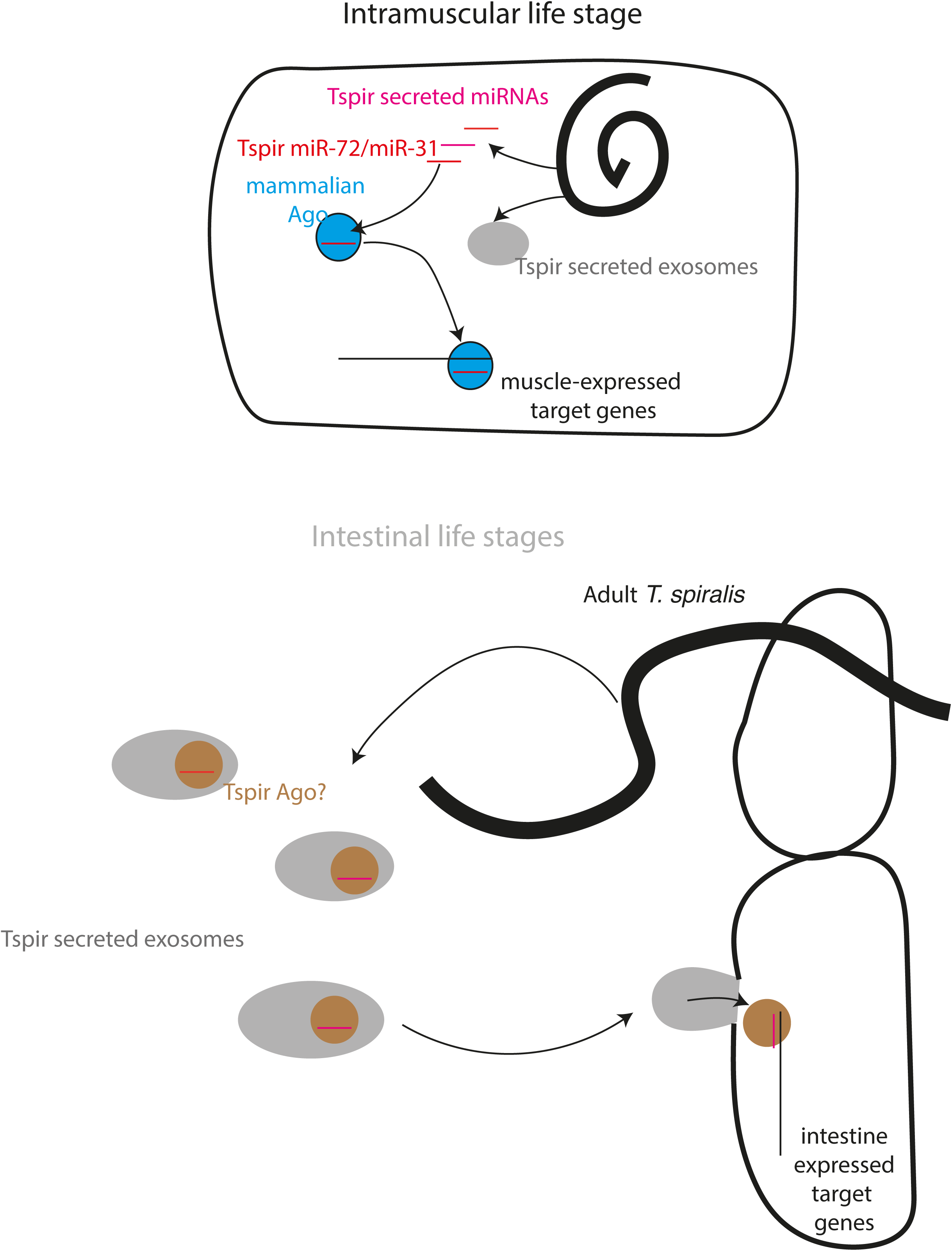
Model showing secretion of small RNAs by MSL *T. spiralis* (top) or Adult *T. spiralis* (bottom) and how these may play roles in host gene expression regulation by the parasite.

The difference between adult and larval *T. spiralis* and apparent secretion of unencapsulated small RNAs by the latter may also relate to the unusual life cycle of this nematode. In the intestinal phase, development of infective larvae to adult worms requires invasion of epithelia (Gagliardo, 2002; ManWarren et al., 1997). Although described as occupying a multicellular niche, worms migrate through epithelial cell monolayers in vitro leaving a trail of dead cells (ManWarren et al., 1997), and have been observed migrating in and out of the epithelial cell layer *in vivo* (Wright, 1979). This is therefore not an intracellular parasite in the conventional sense in terms of permanent enclosure and development within a single cell.

In contrast, invasion of myofibres by first stage larvae and subsequent development is entirely intracellular, forming an interaction which can remain stable for years. The early stages of remodeling of skeletal muscle by *T. spiralis* has gross similarities to repair of muscle following injury with respect to recruitment, activation and proliferation of satellite cells, presumably in response to the damage caused by parasite invasion (Wu et al., 2008). The subsequent processes diverge, leading to repair and regeneration of a contractile myofibre following injury, and remodelling into a nurse cell by *T. spiralis*. In this respect, secretion of a homologue of miR-31 by *T. spiralis* larvae is interesting, given its key role in repression of the myogenic programme (Crist et al., 2012). Moreover, miR-31 expression is associated with Duchenne Muscular Dystrophy (DMD): it is found at higher levels in human DMD biopsies, and its persistent upregulation in mouse models of the disease (mdx mice) is linked to delay of the muscle differentiation programme, reduced fibre maturation and intensive regeneration (Cacchiarelli et al., 2011; Greco et al., 2009).

Intracellular location of larval *T. spiralis* within skeletal muscle would mean that unencapsulated small RNAs may have a higher chance of engaging with mammalian pathways of gene regulation. Differentiation and repair of skeletal muscle is highly complex, and the cellular remodeling effected by *T. spiralis* infection quite unique, and thus in order to test the plausibility of this model further, confirmation of miR-31/72 and other parasite secreted miRNAs within muscle cells in vivo is required.

## Acknowledgements

This study was supported by a BBSRC studentship to PJT (BB/M0111788/1), BBSRC grants to MES (BB/S001085/1) and MB (BB/L023091/1) and a grant from the Medical Research Council to PS (Transgenerational Epigenetic Inheritance and Evolution)

